# Seizing diminishing habitat conservation and restoration opportunities in the Tampa Bay, FL watershed: An urban estuary in recovery

**DOI:** 10.1101/2020.11.16.384404

**Authors:** Gary E Raulerson, Douglas E Robison, Marcus W Beck, Maya C Burke, Thomas F Ries, Justin A Saarinen, Christine M Sciarrino, Edward T Sherwood, David A Tomasko

**Author notes:** Sarasota Bay Estuary Program, Sarasota, Florida, United States of America (David A Tomasko). Corresponding author: (GR).

## Abstract

Habitat restoration efforts should integrate past trends, current status, expected climate change and coastal development impacts, remaining realistic opportunities, and resource management community capabilities. Integrating these concepts, a new target setting approach is being implemented in the Tampa Bay region with broad transferability potential. Past changes, as determined through a three-decade habitat change analysis and over forty years of habitat restoration experience in the region, has informed the new approach. It is also primarily focused on what is possible today and the projected needs for the future, rather than focusing on or attempting to replicate past ecological conditions. Likewise, this new paradigm accounts for persistent local and global stressors – especially watershed development, sea level rise, and climate change. As such, newly established numeric targets are “place-based,” meaning that they attempt to maximize the remaining restoration and conservation “opportunity areas” within the watershed. Lastly, the approach is comprehensive in that targets for the range of critical habitats, from subtidal to uplands, are now defined. This approach represents a general framework for addressing competing interests in planning for habitat restoration that could be applied in other coastal settings where sustainable urbanization practices are desired to co-exist with natural environments.

## Introduction

The health of estuarine systems, coastal habitats, and associated fauna and flora are inextricably linked to land uses and management throughout the watershed [1]. These habitats provide multiple ecosystem services, including wildlife shelter and migratory corridors [1], fisheries production [3], water quality improvement [4,5,6], erosion and flood attenuation [7,8], carbon sequestration [9] and recreation [10]. The Tampa Bay Estuary Program (TBEP) is one of 28 programs administered by the US Environmental Protection Agency (USEPA) under the National Estuary Program (NEP). In recognition of threats to habitats from development and climate change stressors, the TBEP and partners recently created the third iteration of a plan to establish targets and goals for habitat restoration within the Tampa Bay watershed. As an NEP, the program has guided regional environmental restoration initiatives for the estuary since 1991. The methodologies used in the creation of a 2020 Habitat Master Plan Update for Tampa Bay [11] are highly transferable to both coastal and non-coastal systems.

### Development, climate change, and coastal squeeze

Development causes multiple perturbations within any watershed. The accumulated impacts of construction and associated infrastructure remove or substantially modify existing habitats, and can alter hydrology of nearby streams and rivers [12,13]. Given the degree to which the Tampa Bay watershed has been urbanized, the synergistic effects of continued coastal development, future climate change, and sea level rise are primary concerns for maintenance of estuarine and coastal habitat health. Observed and potential adverse effects of climate change and sea level rise on marine and estuarine ecosystems are well-documented [14,15,16]. With regard to estuarine habitats, the primary concerns are that sea level rise is now occurring at such a rapid rate that the landward migration of tidal wetlands in response cannot keep pace; or that the upland slope has already been lost to urban development and hardening, leaving no place for tidal wetland migration [16]. Geological, physical and chemical changes could include alterations in sediment deposition and erosion patterns, micro-topography, and water quality [17,18].

Related changes in habitats in response to climate change (sea level rise and warming) include landward migration of mangroves into salt marshes, upstream migration of salt marshes within tidal tributaries, upland forest migration, and reduced freeze events [19,20,21]. Species dependent upon these habitats will be forced to change distribution patterns or adapt to the new conditions [22]. For example, while black needle rush (*Juncus roemerianus*) can tolerate a wide salinity range [23,24], the largest remaining *J. roemerianus* marshes in Tampa Bay are located in the lower-salinity reaches of tidal rivers and creeks. The greatest extents of those marshes occur in river systems where their upstream extent is constrained by impoundments for public water supplies. Spatial restriction in these hydrologically truncated rivers may make these marshes particularly vulnerable to “pinching out”, as upstream migration in response to sea level rise will be cut off by anthropogenic barriers. Similarly, landward migration of salt barrens (high marsh areas in Tampa Bay) in response to sea level rise will be restricted by the filling and hardening of coastal uplands associated with existing or future urban development.

### Need for a paradigm shift

The synergistic effects of development and climate change diminish the available space for future restoration in urbanizing estuaries, thereby impacting the variety of ecosystem services provided by these habitats and the wildlife they support [25]. Given projected habitat losses without intervention efforts and the limited resources (including time, land, funding, and labor force) available, it is important to appropriately and realistically site restoration projects to increase the likelihood of success and benefits to both natural and human communities [26]. To achieve this objective, a new restoration approach on a broad watershed scale will be implemented.

Previously, a “retrospective” approach to setting habitat protection and restoration targets in Tampa Bay was employed [27,28,29,30]. Under this paradigm, priority was given to restoration activities focused on habitat types, important for a suite of ten estuarine faunal guilds, that were disproportionately lost or degraded compared to a benchmark period. Primary criticisms of this approach included a lack of consideration for future sea level rise and other climate change factors [1], use of expanded and different habitats outside the Tampa Bay watershed by the faunal guilds [28], lack of attention to upland or freshwater wetland habitats, and little recognition of other stressors such as land development trends or actual available space for restoration efforts.

Past approaches for guiding restoration planning have been successfully used in other contexts, but they do not fully balance competing needs. For example, an integrated watershed approach [31] has been utilized since the early 1990s to diagnose and manage water quantity and quality problems that have contributed to seagrass restoration in the system. Additionally, the habitat mosaic approach [32] of including multiple habitat types within restoration projects is recognized as necessary in Tampa Bay [33] and elsewhere to allow for ecosystem state changes in response to different environmental pressures [34,35].

Adaptive management [41,42] components have been increasingly used to address challenges of sea level rise, climate change, and development stressors, including monitoring to identify critical restoration decision points and needed intervention with contingency plans. Rising sea levels and temperatures and altered rainfall patterns are causing observable changes to habitats on a global scale [36,37,1], including within Tampa Bay [38], and those changes are expected to become more pronounced over the next several decades [39,40].

### Updating the approach

The new approach integrates the whole watershed, addresses historical changes, focuses on trajectories that have occurred during more contemporary time periods, and considers both current and future stressors -- particularly land development and sea level rise. There is relatively consistent extent and distribution data for most Tampa Bay habitats of interest (1988 to 2018), representing a time period when federal, state and local regulations were in effect and regional impacts from climate change are documented [43,36]. This approach establishes a broader framework that guides both watershed-level habitat master planning and site-level restoration design activities and incorporates applicable elements of the other habitat restoration paradigms discussed above [35]. Our broader framework for guiding restoration activities includes: 1) designation of habitat types by watershed strata relative to the aquatic-terrestrial gradient (tidal wetlands, contiguous freshwater wetlands, isolated freshwater wetlands, and native uplands); 2) quantification of historical trends by habitat types to identify appropriate future targets in acreage; and 3) identification of opportunity areas that could be used by practitioners to achieve restoration goals based on habitat type and past trajectories.

With regard to habitat restoration projects, the design approach must envision not only what is possible today, but also what the coastal landscape will look like in 50 years and beyond. Design features should continue to use the historical “habitat mosaic” approach, but should also include coastal upland features that accommodate tidal inundation and the landward advance of emergent tidal wetlands [44].

## Methods

### Study area

Tampa Bay is a large open water estuary (open water area approximately 983 km^2^) on the west-central coast of Florida (Fig 1). Its watershed encompasses another 5,872 km^2^, for a total combined area of approximately 6,855 km^2^. It is subtropical, and within the current (2020) ecotone for mangrove and salt marsh habitats [45]. The upstream watershed includes multiple habitats, including pine flatwoods, forested freshwater wetlands and non-forested vegetated wetlands. The watershed is heavily developed with an estimated (2019) population of 3.3 million people in the four counties that comprise most of the watershed [46]. Numerous anthropogenic changes have been made to the natural systems within and surrounding Tampa Bay, including direct removal of habitat (including dredge and fill of bay bottom), alteration of hydrology, and destruction and fragmentation of habitat from development.

**Fig 1.**
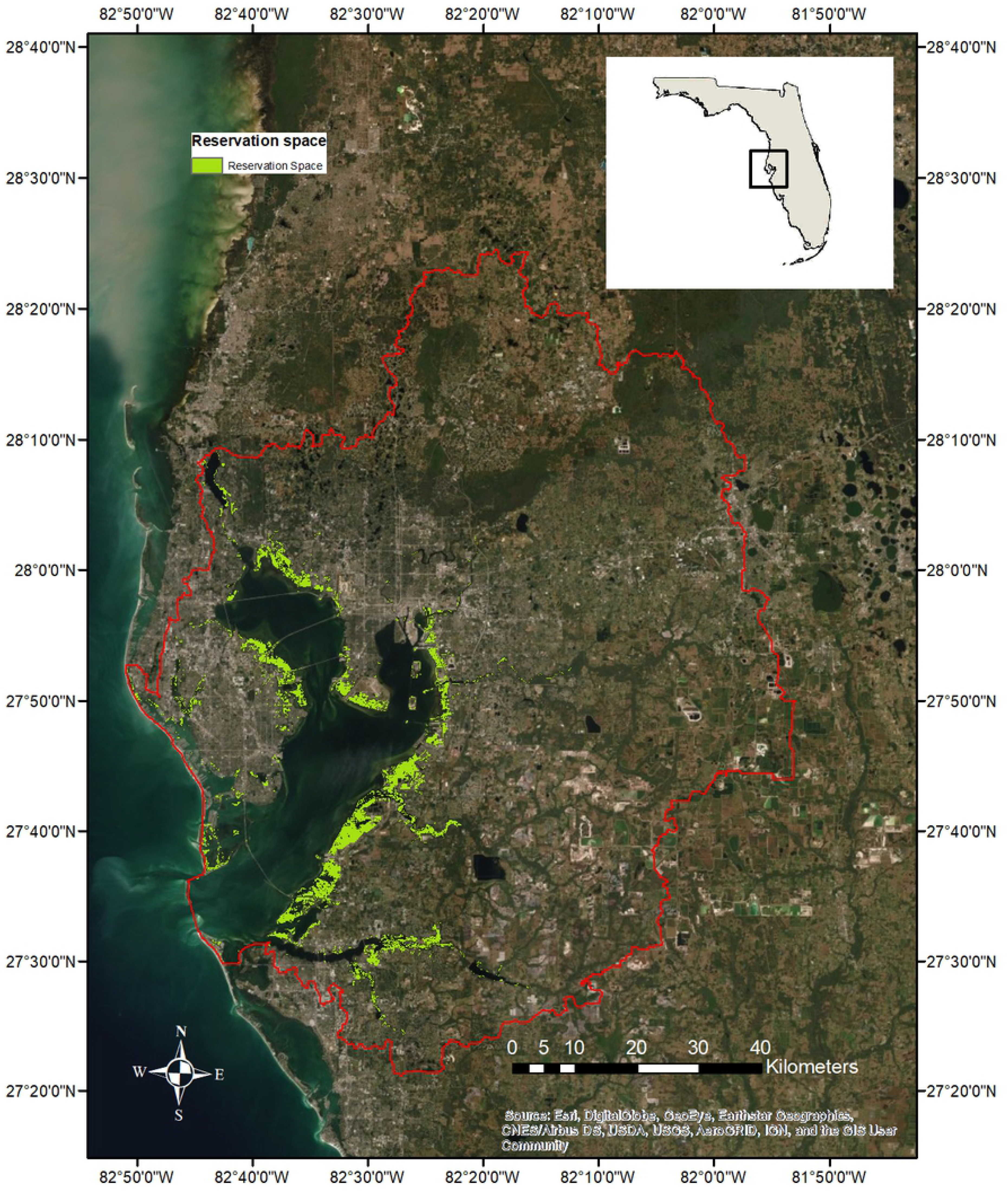
Tampa Bay watershed and defined coastal reservation space.

### Habitats of Tampa Bay

Addressing the full suite of habitats in a watershed is now recognized as critical for large-scale restoration planning efforts [35,47]. The major habitat types of Tampa Bay can be described and organized relative to tidal influence and location in the watershed. Subtidal habitats include those that are submerged all or most of the time; emergent tidal wetlands include those that are submerged during high tides but exposed during low tides; and supratidal habitats include those that occur above the high tide line.

Habitats generally described as ‘subtidal’ include hard bottom [48,49], artificial reefs [50], tidal flats [51,52], seagrasses [53,54], and oyster reefs [55,56]. Mangroves [57], salt marshes [43,58], salt barrens [59,60], tidal tributaries [61,62,63,64], and living shorelines [65,66,67] are classified as emergent tidal wetlands. For the purposes of this planning effort, supratidal habitats included non-developed uplands [45], freshwater forested wetlands [68], and freshwater non-forested wetlands [4]. As discussed below, uplands are sub-divided into coastal and non-coastal uplands, based on location relative to the 5-foot contour.

### Approach

Recommended targets were created stepwise with broad-scale suitability modeling within a GIS environment: 1) quantifying habitat status and historical trends; 2) synthesizing historic habitat restoration efforts; and 3) identifying and defining remaining ‘opportunities’ for restoration that integrated results from the first two analyses. Final development of restoration targets and goals integrated strata and opportunity area information with an assessment of decadal-scale restoration investments as a guide to define realistic and practical future restoration to be conducted by partners. This approach further supported estuarine-dependent species and faunal guilds throughout the watershed, as defined in the original approach, while also framing future restoration investments in the context of the practical pace employed by the region during the Bay’s initial recovery.

### Habitat status and trends

For the majority of subtidal, intertidal and supratidal habitats, primary data derived from two routine spatial assessment programs conducted by the Southwest Florida Water Management District (SWFWMD) were utilized. However, to address data gaps for some habitats, results from special studies were integrated with these primary data sources. These included targeted studies to better understand hard bottom [69,70], dredged holes [71,72] and oyster habitat [73] within Tampa Bay.

The source data used to estimate the most current coverage of seagrasses, tidal flats, and oysters was the *Seagrass in 2018* geospatial database [74]. The bi-annual seagrass monitoring program was initiated in 1988 under SWFWMD’s Surface Water Improvement and Management program [54,75]. SWFWMD has estimated oyster bed coverage as part of this program since 2014.

The source data used to estimate and map trends in development, emergent tidal wetlands, freshwater wetlands, and native upland habitats, was the SWFWMD Land Use Land Cover series geospatial database [76]. This comprehensive database classifies the land use and cover types (natural and developed) pursuant to the Florida Land Use Cover and Forms Classification System (FLUCCS)[77,78]. Mangroves, salt barrens, and salt marshes were reported individually. While the photointerpretation of specific freshwater wetland types is often very difficult, it is possible to accurately distinguish forested wetlands from non-forested wetlands. Therefore, for this analysis, all applicable FLUCCS codes representing the suite of natural freshwater wetlands were combined within those two classifications. Similarly, within the target- and goal-setting exercise, uplands were combined in one classification. These classifications were reported for each mapping exercise conducted every 2-3 years from the start of the program 1990 through 2017, the most recent year with available data.

To quantify the extent of tidal creeks, FLUCCS datasets [62,63] were extracted to the Florida Department of Environmental Protection stream segments that were classified as estuarine.

### Restoration database

We quantified past restoration effort in each of the major habitat types to guide decisions on future targets and goals. Information regarding habitat restoration and enhancement activities in the Tampa Bay area over the past 40 years were compiled, reviewed, and consolidated into a single, consistent geospatial database. Data were gathered from the SWFWMD Surface Water Improvement and Management Program, Federal Government Performance and Results Act reporting, the Tampa Bay Water Atlas (https://www.tampabay.wateratlas.usf.edu/), restoration practitioners (e.g. the non-profit Tampa Bay Watch) and the Technical Advisory Committee of the TBEP. Primary data collected included project name, year, description, lead partner, size (area or length), and latitude and longitude. Data gaps were supplemented by archival research, site visits, contacting the responsible entities, and documenting knowledge of local restoration practitioners. Living shoreline projects, including seawall enhancements and oyster reef modules, were inventoried separately.

### Creating and combining opportunity layers

Three distinct geographic breakpoints, or strata, in the watershed where different habitat management and restoration activities could take place, were created to establish the process for target and goal-setting within this framework. The first is the coastal, or reservation stratum (Table 1), extended from the local Mean Lower Low Water elevation to elevation 5 feet above Mean Sea Level and is likely to be affected by frequent tidal flooding or inundation by 2070. The reservation stratum is the zone where emergent tidal wetland restoration would be conducted. It includes low lying coastal uplands which serve as important tidal wetland buffers that will be critically important in the future as lands reserved to accommodate ideal wetland migration in response to sea level rise. The river floodplain stratum includes all hydrologically contiguous forested and non-forested wetlands within the river and stream corridors of the Tampa Bay watershed. Floodplain corridors provide vital watershed functions including fish and wildlife habitat and migratory pathways, floodwater attenuation and storage, erosion control, and delivery of complex organic matter to the estuarine food web. Finally, the upland stratum encompasses those areas outside of the coastal and river floodplain strata, including native upland habitats as well as hydrologically isolated wetlands. These habitats provide important aquifer recharge and wildlife habitat functions.

**Table 1.**
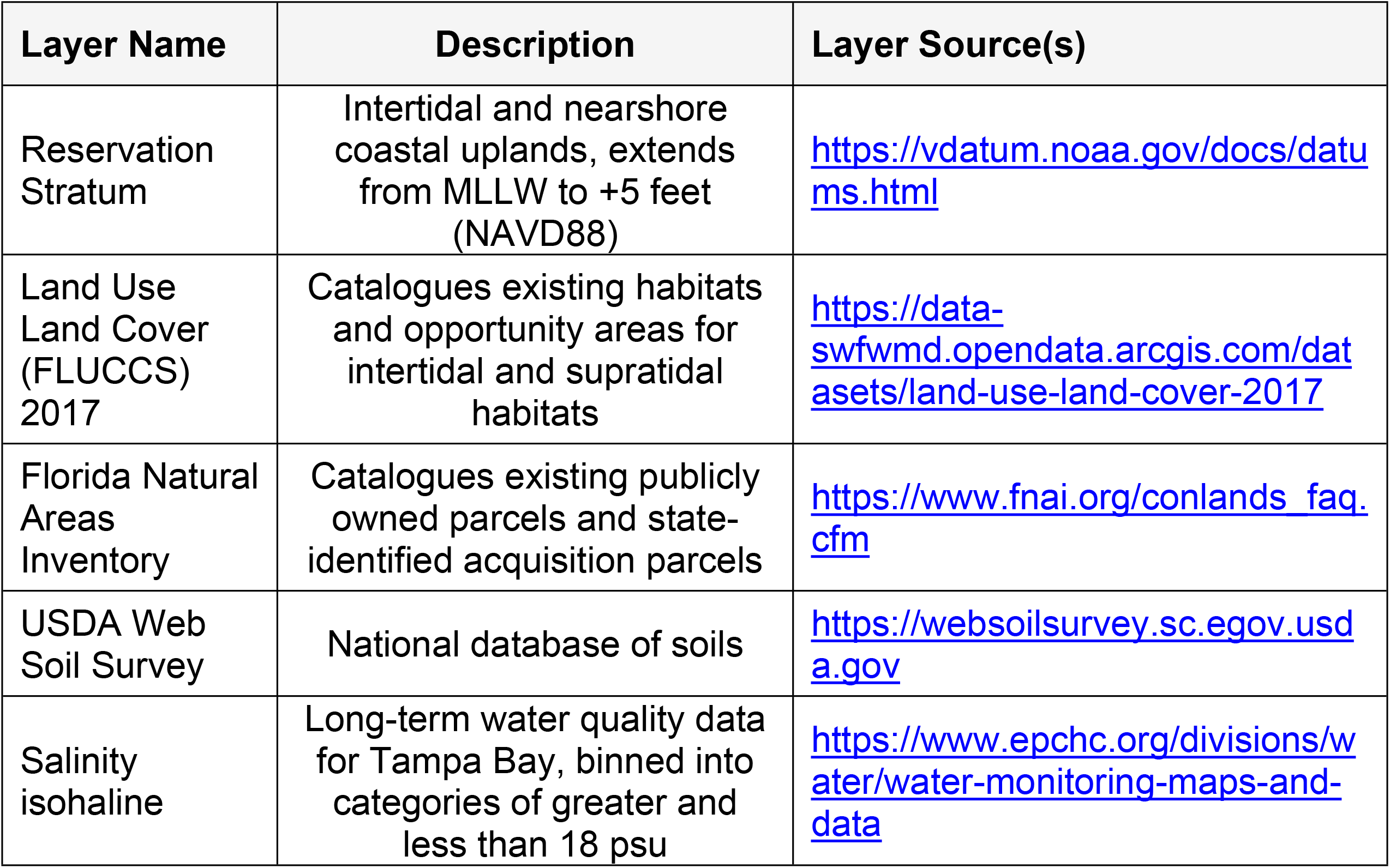
GIS layers used to identify opportunity areas for restoration targets and goals.

Opportunity areas, defined here as locations where habitat protection and restoration activities are possible, and where they should best be focused to attain defined targets in the future, were also analyzed. The definition and mapping of opportunity areas is necessary to quantify the “restoration potential” for a particular habitat type, which is a measure of what is actually possible under current and future projected conditions. The most appropriate opportunity areas were generally not developed and located on existing public lands or areas identified for public acquisition as conservation lands.

The FLUCCS 2017 geospatial database (Table 1) was used as the baseline for cataloguing existing and opportunity areas for intertidal and supratidal habitats in the Tampa Bay watershed. All FLUCCS classification codes were placed into one of three categories. First, native habitats cover the full range of natural plant communities and other habitats that are endemic to the Tampa Bay watershed, and were further grouped into three major habitat types (tidal wetlands, freshwater wetlands, and uplands). Second, restorable habitats include existing altered but non-hardened and pervious FLUCCS codes that could potentially support native habitats through the restoration of more natural hydrology, soils strata, and/or topography. Third, existing development includes developed land FLUCCS codes that are hardened and impervious (e.g., structures and pavement) and difficult to prioritize for habitat restoration activities.

Layers for existing public lands and parcels targeted for acquisition were compiled by combining data from the Florida Natural Areas Inventory (Table 1), consulting staff from various federal, state and local entities, and inventorying official conservation and drainage easement records data.

Compared to vegetation communities, soil characteristics typically change slowly (e.g., decades to centuries) in response to hydrologic impacts, unless physically disturbed [79,80]. Therefore, soils distributions can be used to generally represent historical habitat distributions, and can be used to provide generalized restoration guidelines (e.g., tidal wetlands, freshwater wetlands, and native uplands). Ries and Scheda [81] created a soils suitability analysis for wetland mitigation and restoration using data from the USDA Web Soil Survey (Table 1) and classified all soils in the Tampa Bay watershed into one of three categories (xeric, mesic, and hydric). The mesic and hydric categories were combined to represent wetland restoration potential, while the xeric category was used to represent upland restoration potential.

A distinction was made between tidal and freshwater wetland restoration potential by intersecting the combined mesic and hydric soils polygons with the coastal stratum. Mesic or hydric soils that occur below the 5-foot contour were classified as having tidal wetland restoration potential, while mesic or hydric soils occurring above the 5-foot contour were classified as having freshwater wetland restoration potential.

Within the Tampa Bay region, lands adjacent to waters with average salinity values greater than 18 psu were considered most appropriate for higher salinity mangrove/salt barren restoration, while lands adjacent to waters with average salinity values less than 18 psu were considered most appropriate for lower salinity salt marsh (*Juncus* spp.) restoration [82]. To estimate the relative restoration potential of mangrove/salt barrens and salt marshes, a regional long-term water quality data set was used to create salinity isohalines, which was then binned into two salinity categories: greater and less than an annual mean of 18 psu (Table 1).

All source and derivative GIS datasets (Table 1) were converted into uniform 10m x 10m raster datasets extracted to the Tampa Bay watershed boundary using spatial analyst tools in ArcGIS Pro 2.x. FLUCCS data were reclassified into appropriate restoration categories (native habitats, non-native habitats and non-restorable) and combined with the reservation, restorable, conservation, acquisition, soils, and salinity isohaline layers to create a comprehensive index for potential restoration opportunity areas identified as existing and proposed restoration areas, reservation areas, soil type, and salinity level.

## Results

### Habitat status and trends

#### Subtidal habitats

While no trend information is available, the best cumulative estimate of natural hard bottom extent in Tampa Bay is 171 ha (Table 2). Oyster bars covered 69 ha (Table 2) in 2018, and a 30% (6 ha) increase was reflected since mapping began in 2014 (Table 3a). This increase probably represents improved ground-truthing and photointerpretation of oyster bar signatures from aerial photography, rather than actual expansion of oyster habitat. Twelve artificial reefs in Tampa Bay are managed by Hillsborough, Manatee, and Pinellas Counties. Surface area estimates were not available for the Manatee and Pinellas County reefs, but assuming an average size of 4.2 ha, based on the Hillsborough County reefs, the total coverage of artificial reefs in Tampa Bay is estimated to be approximately 67 ha (Table 2).

**Table 2.** Current extent and data sources for Tampa Bay habitats.

| Habitat Type | Current Extent | Data Year | Data Source(s) |
| --- | --- | --- | --- |
| **Subtidal Habitats** |  |  |  |
| Hard Bottom | 171 ha | 2017 - 2019 | SWFWMD, 2017; CSA Ocean Sciences, 2019 |
| Artificial Reefs | 67 ha | 2019 | EPCHC, 2020; ESA, 2020 |
| Tidal Flats | 6,564 ha | 2018 | SWFWMD, 2018 |
| Seagrasses | 16,452 ha | 2018 | SWFWMD, 2018 |
| Oyster Bars | 69 ha | 2018 | SWFWMD, 2018 |
| **Intertidal Habitats** |  |  |  |
| Living Shorelines | 18.2 km | 2020 | ESA, 2020; Tampa Bay Watch, 2020 |
| Mangrove Forests | 6,192 ha | 2017 | SWFWMD, 2017 |
| Salt Barrens | 201 ha | 2017 | SWFWMD, 2017 |
| Salt Marshes | 1,844 ha | 2017 | SWFWMD, 2017 |
| Tidal Tributaries | 623 km | 2019 | Janicki Environmental/Mote Marine Lab, 2019 |
| **Supratidal Habitats** |  |  |  |
| Coastal Uplands | 1,465 ha | 2017 | ESA, 2020 |
| Non-Forested Freshwater Wetlands | 27,352 ha | 2017 | SWFWMD land use/cover mapping |
| Forested Freshwater Wetlands | 61,566 ha | 2017 | SWFWMD land use/cover mapping |
| Native Uplands (Non-Coastal) | 56,899 ha | 2017 | SWFWMD land use/cover mapping |

**Table 3a.** Subtidal habitat trends in Tampa Bay, 1988-2018.

| Habitat Descriptor | 1988 | 1990 | 1992 | 1994 | 1996 | 1999 | 2001 | 2004 | 2006 | 2008 | 2010 | 2012 | 2014 | 2016 | 2018 | 1988-2018 Change |  |
| --- | --- | --- | --- | --- | --- | --- | --- | --- | --- | --- | --- | --- | --- | --- | --- | --- | --- |
|  |  |  |  |  |  |  |  |  |  |  |  |  |  |  |  | Hectares | Percent |
| Seagrass Total | 9,424 | 10,210 | 10,424 | 10,736 | 10,901 | 10,460 | 10,555 | 10,938 | 11,453 | 11,997 | 13,313 | 14,020 | 16,307 | 16,858 | 16,452 | 7,027 | 75% |
| Non-Vegetated Total | 11,084 | 10,367 | 10,561 | 10,305 | 10,492 | 13,231 | 12,642 | 14,631 | 14,684 | 13,473 | 11,649 | 10,360 | 7,106 | 5,711 | 6,564 | -4,520 | -55% |
| Oysters Total | ND | ND | ND | ND | ND | ND | ND | ND | ND | ND | ND | ND | 53 | 68 | 69 | NA | NA |

Total seagrass coverage has increased by 7,027 ha (75%) during the 30-year period of record (Table 3a), and the most current (2018) estimate of total seagrass meadow coverage in Tampa Bay is 16,452 ha. The 2018 coverage of tidal flats and sand other than beaches in Tampa Bay was 6,564 ha (Table 2). A decrease of 4,496 ha (55%) during the 30-year period of record (Table 3a) is partly associated with the expansion of seagrass to previously non-vegetated bottom area, as well as changes in map classification protocols.

#### Intertidal habitats

Between 1990 and 2017, the suite of emergent tidal wetlands (mangroves, salt barrens, and salt marshes) experienced a net gain of 725 ha (10%, Table 3b). The current estimate of mangrove forest extent in Tampa Bay is 6,192 ha. Mangrove forest coverage increased by 684 ha (12%). Salt marshes in Tampa Bay cover 1,844 ha, and coverage increased by 30 ha (2%). However, from 1990 to 2017, it is estimated that a net area of 219 ha of salt marshes converted to mangrove habitat [83]. The 2017 estimate of the extent of salt barrens in Tampa Bay is 201 ha, and coverage has increased by 14 ha (7%) during the 27-year period of record (Table 3b). Based on GIS data from Janicki Environmental and Mote Marine Laboratory [62,63], the extent of tidal creek habitat in the Tampa Bay watershed is approximately 623 km (Table 2). No trend analysis is available.

**Table 3b.**
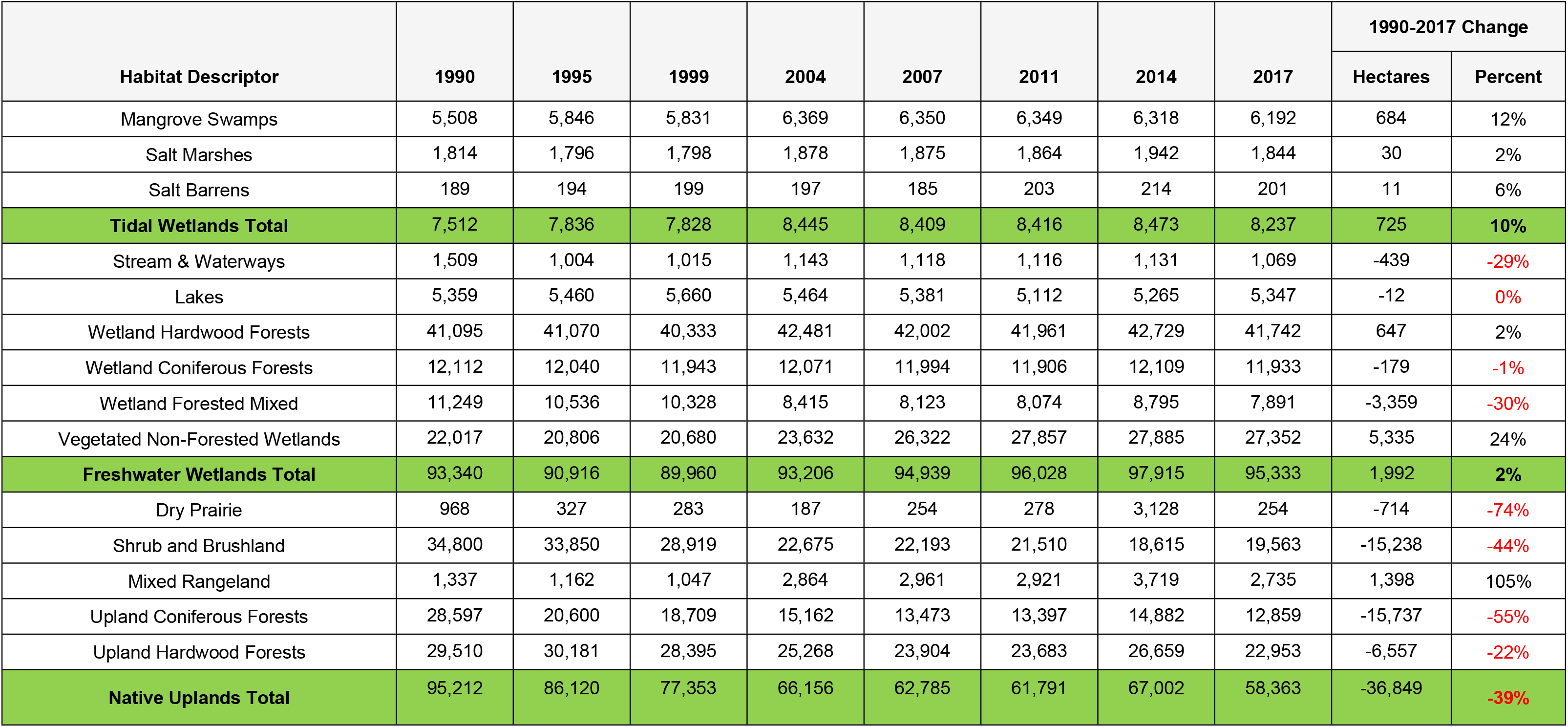
Intertidal and supratidal habitat trends in Tampa Bay, 1990-2017.

| Habitat Descriptor | 1990 | 1995 | 1999 | 2004 | 2007 | 2011 | 2014 | 2017 | 1990-2017 Change |  |
| --- | --- | --- | --- | --- | --- | --- | --- | --- | --- | --- |
|  |  |  |  |  |  |  |  |  | Hectares | Percent |
| Mangrove Swamps | 5,508 | 5,846 | 5,831 | 6,369 | 6,350 | 6,349 | 6,318 | 6,192 | 684 | 12% |
| Salt Marshes | 1,814 | 1,796 | 1,798 | 1,878 | 1,875 | 1,864 | 1,942 | 1,844 | 30 | 2% |
| Salt Barrens | 189 | 194 | 199 | 197 | 185 | 203 | 214 | 201 | 11 | 6% |
| **Tidal Wetlands Total** | **7,512** | **7,836** | **7,828** | **8,445** | **8,409** | **8,416** | **8,473** | **8,237** | **725** | **10%** |
| Stream & Waterways | 1,509 | 1,004 | 1,015 | 1,143 | 1,118 | 1,116 | 1,131 | 1,069 | -439 | -29% |
| Lakes | 5,359 | 5,460 | 5,660 | 5,464 | 5,381 | 5,112 | 5,265 | 5,347 | -12 | 0% |
| Wetland Hardwood Forests | 41,095 | 41,070 | 40,333 | 42,481 | 42,002 | 41,961 | 42,729 | 41,742 | 647 | 2% |
| Wetland Coniferous Forests | 12,112 | 12,040 | 11,943 | 12,071 | 11,994 | 11,906 | 12,109 | 11,933 | -179 | -1% |
| Wetland Forested Mixed | 11,249 | 10,536 | 10,328 | 8,415 | 8,123 | 8,074 | 8,795 | 7,891 | -3,359 | -30% |
| Vegetated Non-Forested Wetlands | 22,017 | 20,806 | 20,680 | 23,632 | 26,322 | 27,857 | 27,885 | 27,352 | 5,335 | 24% |
| **Freshwater Wetlands Total** | **93,340** | **90,916** | **89,960** | **93,206** | **94,939** | **96,028** | **97,915** | **95,333** | **1,992** | **2%** |
| Dry Prairie | 968 | 327 | 283 | 187 | 254 | 278 | 3,128 | 254 | -714 | -74% |
| Shrub and Brushland | 34,800 | 33,850 | 28,919 | 22,675 | 22,193 | 21,510 | 18,615 | 19,563 | -15,238 | -44% |
| Mixed Rangeland | 1,337 | 1,162 | 1,047 | 2,864 | 2,961 | 2,921 | 3,719 | 2,735 | 1,398 | 105% |
| Upland Coniferous Forests | 28,597 | 20,600 | 18,709 | 15,162 | 13,473 | 13,397 | 14,882 | 12,859 | -15,737 | -55% |
| Upland Hardwood Forests | 29,510 | 30,181 | 28,395 | 25,268 | 23,904 | 23,683 | 26,659 | 22,953 | -6,557 | -22% |
| **Native Uplands Total** | **95,212** | **86,120** | **77,353** | **66,156** | **62,785** | **61,791** | **67,002** | **58,363** | **-36,849** | **-39%** |

#### Supratidal habitats

The 2017 extent of freshwater wetlands in the Tampa Bay watershed was 88,917 ha (Table 2). Of this total, forested freshwater wetlands comprised 61,565 ha (69%), while non-forested freshwater wetlands comprised 27,351 ha (31%). From 1990-2017, the suite of freshwater wetlands experienced a net gain of 2,444 ha (3%, Table 3b). There has been a 5,335 ha (24%) increase in vegetated non-forested freshwater wetlands since 1990, while forested freshwater wetlands have decreased by 2,891 ha (4%).

As determined in the 2017 land use/land cover update, the most current estimate of the extent of non-coastal native upland habitats in the Tampa Bay watershed is 56,899 ha, and the extent of “coastal uplands” (defined as below the 5-foot contour) in the Tampa Bay watershed is 1,465 ha (Table 2). Over the 27-year period of record, the suite of native upland habitats has experienced a net loss of 37,051 ha (39%, Table 3b).

### Habitat restoration

A total of 460 projects were documented between 1971 and 2019 (Table 4), addressing the full range of habitat types, including Estuarine (n=228), Freshwater (n=53), Uplands (n=119), and Mixed (n=60). A total of 1,978 ha have been restored, and 12,930 ha and 42.8 km of linear projects were enhanced during the time period. Forty lead partners were documented as responsible for the projects, although some of these lead partners are departments within the same agency. Eighty-nine living shoreline projects, seawall enhancements, and oyster reef module installations along shorelines were inventoried, with a linear footprint of 18.2 km.

**Table 4.**
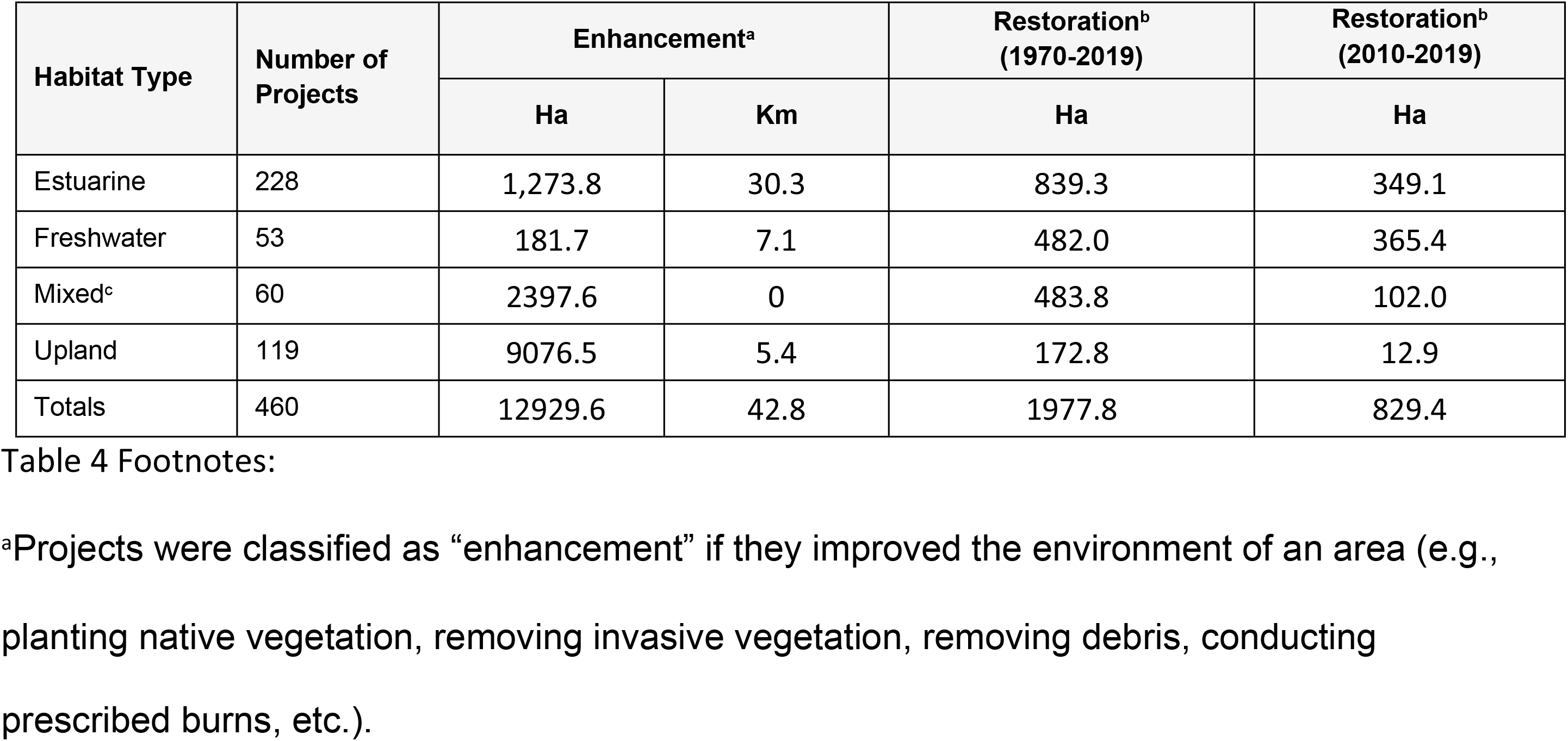
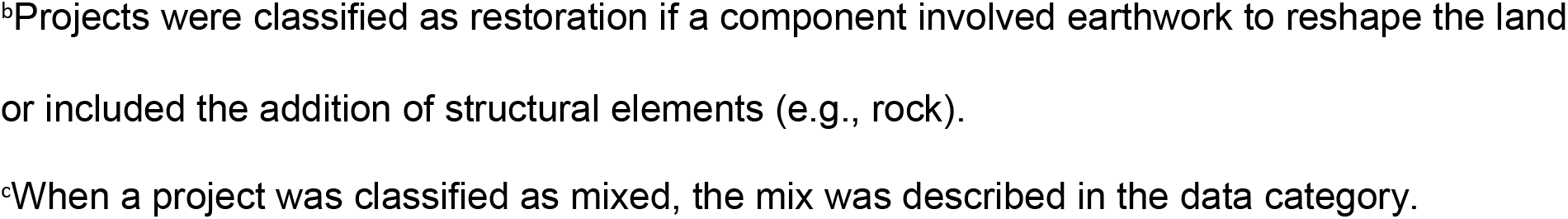
Enhancement and Restoration Projects in Tampa Bay, 1970-2019.

| Habitat Type | Number of Projects | Enhancement<sup>a</sup> |  | Restoration<sup>b</sup><br>(1970-2019) | Restoration<sup>b</sup><br>(2010-2019) |
| --- | --- | --- | --- | --- | --- |
|  |  | Ha | Km | Ha | Ha |
| Estuarine | 228 | 1,273.8 | 30.3 | 839.3 | 349.1 |
| Freshwater | 53 | 181.7 | 7.1 | 482.0 | 365.4 |
| Mixed<sup>c</sup> | 60 | 2397.6 | 0 | 483.8 | 102.0 |
| Upland | 119 | 9076.5 | 5.4 | 172.8 | 12.9 |
| Totals | 460 | 12929.6 | 42.8 | 1977.8 | 829.4 |

<sup>a</sup>Projects were classified as “enhancement” if they improved the environment of an area (e.g., planting native vegetation, removing invasive vegetation, removing debris, conducting prescribed burns, etc.).
<sup>b</sup>Projects were classified as restoration if a component involved earthwork to reshape the land or included the addition of structural elements (e.g., rock).
<sup>c</sup>When a project was classified as mixed, the mix was described in the data category.

### Habitat restoration opportunities

In 2017, 1,555 km^2^ of land (26.5%) in the Tampa Bay watershed above the MLLW line was classified as natural and 2,144 km^2^ (37%) was considered restorable. Developed areas in the Tampa Bay watershed encompassed 2,172 km^2^ (36.5%) of the 5,872 km^2^ watershed area. Between 2014 and 2017, the developed footprint increased by seven percent (7%, Fig 2).

**Fig 2.**
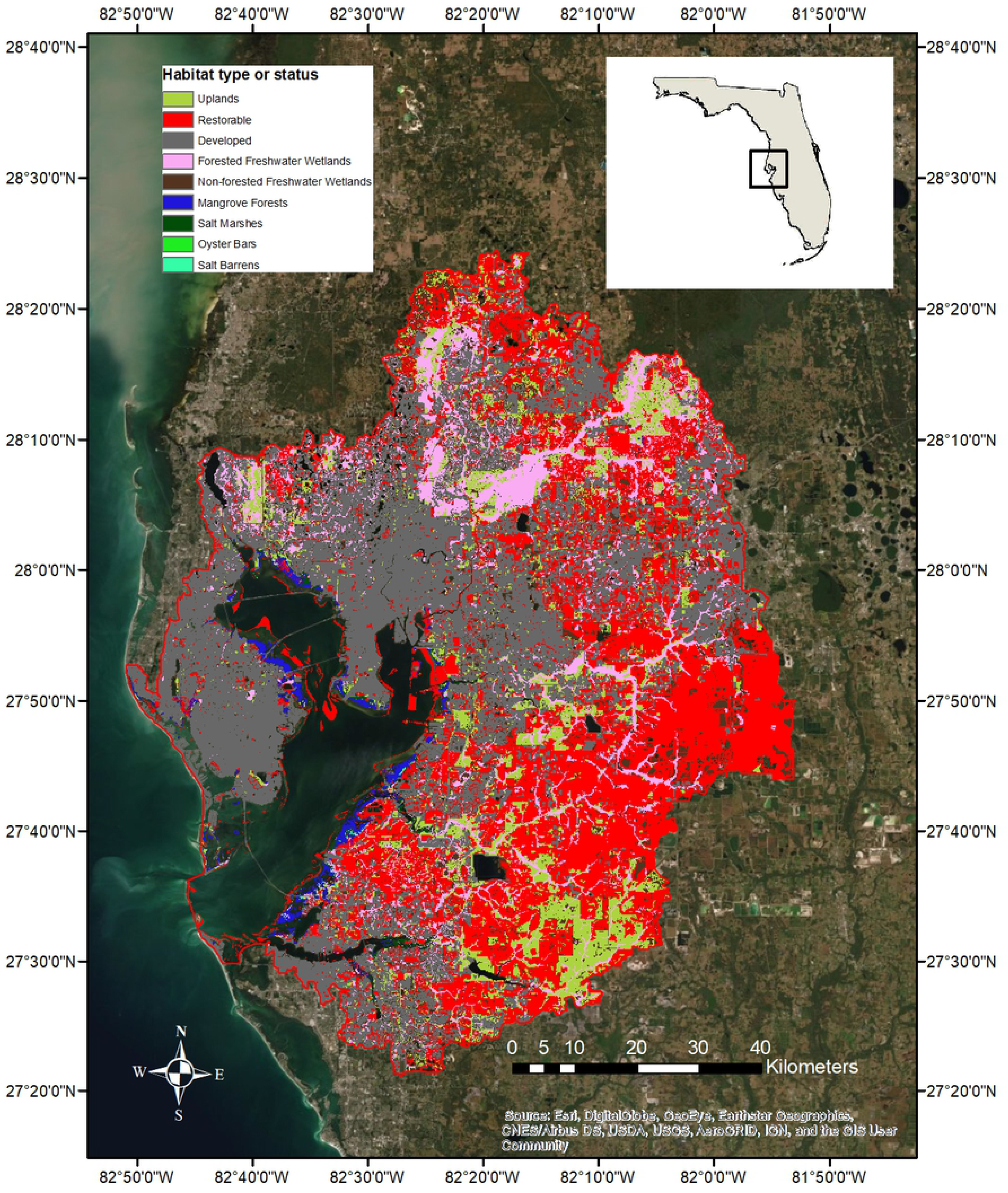
Natural habitats and restorable and developed areas within the Tampa Bay watershed.

The Tampa Bay watershed included a total of 1,260 km^2^ of existing conservation lands, either publicly owned or in conservation easements. However, when excluding subtidal areas owned by the State of Florida, a total of 816 km^2^ of conservation lands is present, about 13.9% of the watershed area occurring above the MLLW line. (Fig 3). Proposed conservation lands in the Tampa Bay watershed total 1,254 km^2^. The mapped proposed conservation lands (Fig 3) generally link eastward and provide wildlife connectivity to broader state conservation initiatives and wildlife corridors [84,85].

**Fig 3.**
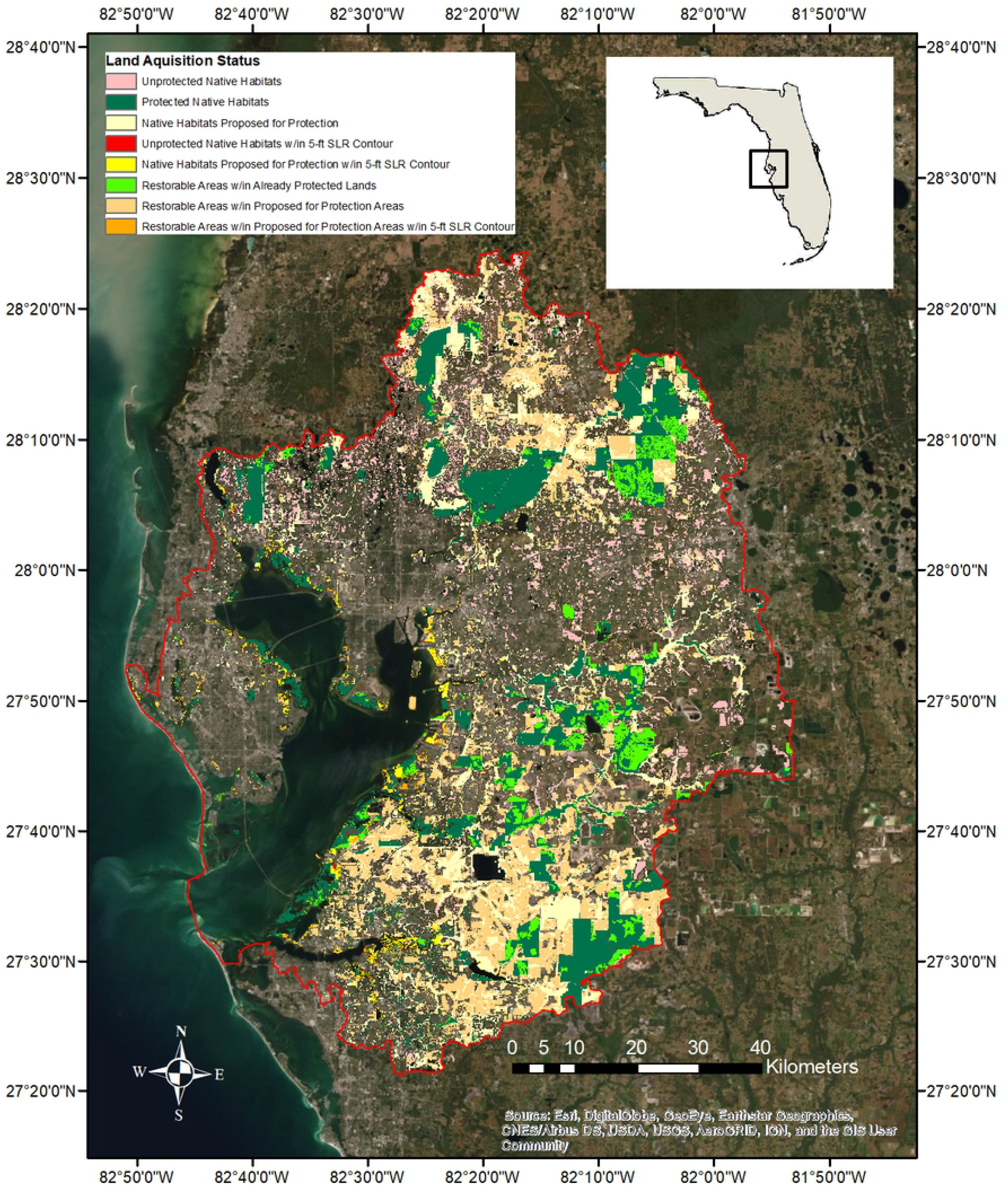
Existing conservation lands and areas proposed for acquisition within the Tampa Bay watershed.

Reservation lands, from the MLLW line landward to elevation 5-feet (NAVD 88), currently (2017) include 4,622 ha proposed for conservation in the Tampa Bay watershed, with 3,303 ha of native habitats and 1,319 ha of restorable habitats (Fig 1).

Xeric soils were roughly aggregated in an east-west band through the middle of the watershed (Fig 4). Approximately 2,107 km^2^ was classified as xeric, while 3,760 km^2^ was classified as mesic/hydric.

**Fig 4.**
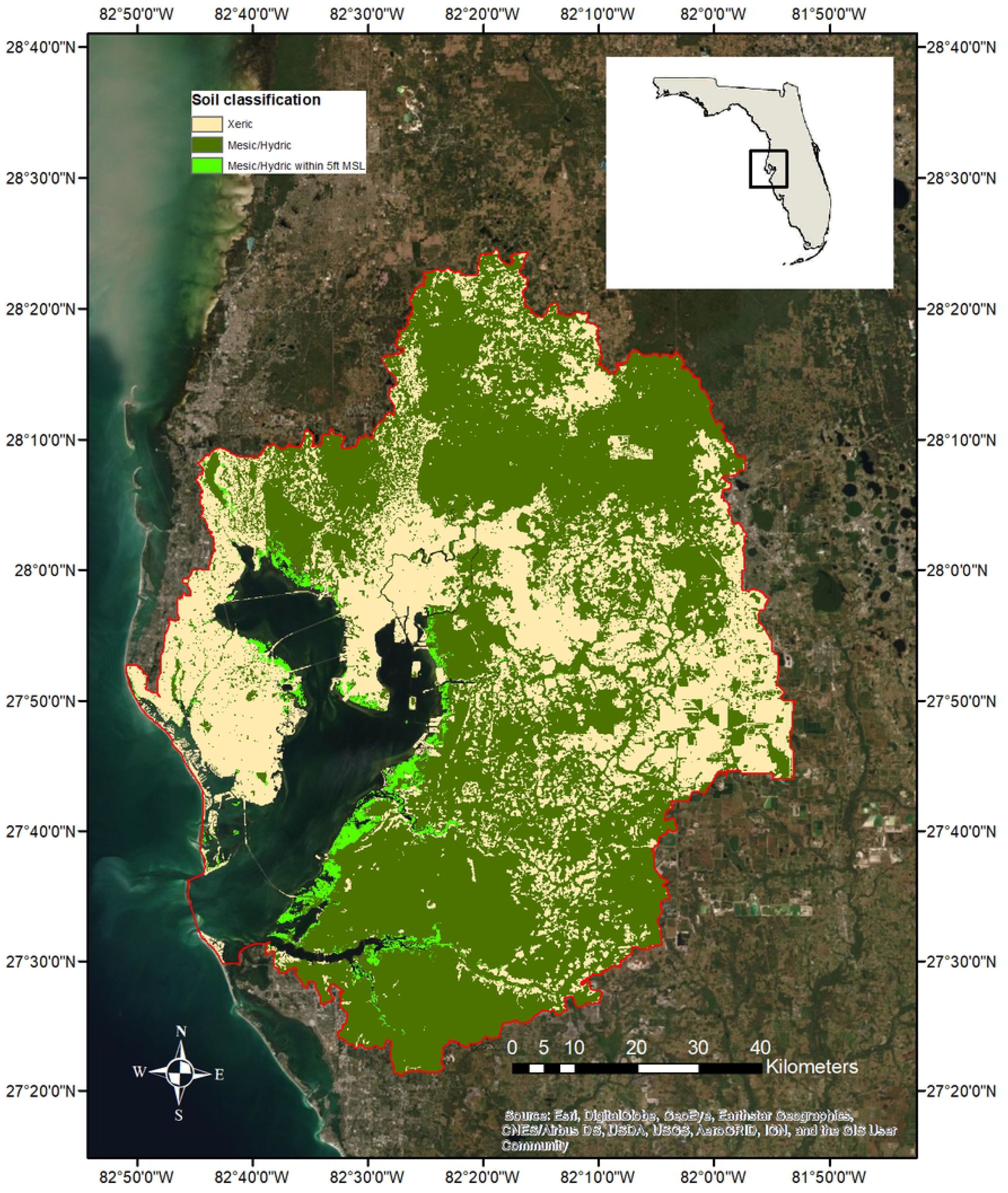
Xeric, mesic, and hydric soils within the Tampa Bay watershed.

Integration of the opportunity layers provides a summary of the available restoration area for the habitats of interest within the Tampa Bay watershed (Table 5). The “Native Habitats” columns show the total current extent as well as the portion of the current extent occurring on existing conservation lands and proposed conservation lands, respectively. The “Restorable Habitats” columns show the total restoration opportunity area as well as the portion of the total restoration opportunity area on existing and proposed conservation lands. The majority of restoration opportunities on existing conservation lands are for native uplands and freshwater wetlands (Table 5). However, there are approximately 627 ha of emergent tidal wetland restoration opportunities on existing conservation lands, including 530 ha applicable to higher salinity mangrove forests and salt barrens, and 16 ha applicable to lower salinity salt marsh (e.g., *Juncus roemerianus*) restoration and creation.

**Table 5.** Habitat restoration opportunities in the Tampa Bay watershed.

Legend: N/A – Not Applicable; I/D – Insufficient Data; LSSM – Living Shoreline
| Habitat Type | Native Habitats |  |  | Restorable Habitats |  |  |
| --- | --- | --- | --- | --- | --- | --- |
|  | Current Extent | Existing Conservation Lands | Proposed Conservation Lands<sup>a</sup> | Total Restoration Opportunity<sup>a</sup> | Existing Conservation Lands Restoration Opportunity | Proposed Conservation Lands Restoration Opportunity<sup>b</sup> |
| **Subtidal Habitats** |  |  |  |  |  |  |
| Hard Bottom (ha) | 171 | 171 | N/A | N/A | N/A | N/A |
| Artificial Reefs (ha) | 67 | 67 | N/A | N/A | N/A | N/A |
| Tidal Flats (ha) | 6,564 | 6,564 | N/A | I/D | I/D | N/A |
| Seagrasses (ha) | 16,452 | 16,452 | N/A | 5,719 | 5,719 | N/A |
| Oyster Bars (ha) | 69 | 69 | N/A | I/D | I/D | N/A |
| **Intertidal Habitats** |  |  |  |  |  |  |
| Living Shorelines (km) | 18 | LSSM | N/A | LSSM | N/A | N/A |
| Mangrove Forests (ha) | 6,192 | 4,397 | 1,650 | 1,116 | 530 | 586 |
| Salt Barrens (ha) | 201 | 174 | 33 |  |  |  |
| Salt Marshes (low salinity *Juncus*) (ha) | 1,844 | 851 | 937 | 442 | 98 | 344 |
| Tidal Tributaries (km) | 623 | N/A | N/A | LSSM | N/A | N/A |
| **Supratidal Habitats** |  |  |  |  |  |  |
| Coastal Uplands (ha) | 1,465 | 698 | 690 | 515 | 126 | 389 |
| Non-Forested Freshwater Wetlands (ha) | 27,352 | 4,647 | 10,510 | 64,683 | 11,107 | 53,576 |
| Forested Freshwater Wetlands (ha) | 61,566 | 23,562 | 22,867 |  |  |  |
| Native Uplands (Non-Coastal) (ha) | 56,899 | 26,051 | 21,381 | 17,777 | 5,368 | 12,409 |

Suitability Model; JU – Potential *Juncus* Marsh Opportunity
<sup>a</sup>All lands identified for acquisition by partners, does not represent a 2030 target or 2050 goal
<sup>b</sup>Does not account for lands neither currently protected nor currently under consideration for acquisition

The best estimates of total restoration opportunities for urban shorelines and tidal tributaries are provided by the Tampa Bay Living Shoreline Suitability Model (LSSM) prepared by the Florida Fish and Wildlife Conservation Commission [84]. There is 2,566 km of shoreline in Tampa Bay, and approximately 33% is recommended for living shoreline enhancement.

### Establishment of goals and targets

Recommended targets were based on habitat status and trends, habitat restoration data, identified restoration opportunities, and current and anticipated trends in development, available funding, and regulations [87,88].

The targets and goals (Table 6) identify where regional consensus for thriving habitats and abundant wildlife could be implemented (as defined in the TBEP 2017 CCMP Update and the 2021-2025 Strategic Plan [88]). However, establishment of these benchmarks also recognizes that the identified habitat protection and restoration areas will change over time, and require that analyses are revisited on 10-year recurring cycles. Further, the 30-year planning horizon (2050) and goals were also identified to be consistent with regional sea level rise projections developed specifically for Tampa Bay [90]. The coastal stratum (from the existing mean low water line to the approximate 5-foot contour) is projected to directly experience the effects of sea level rise by 2050, and is the primary focus area for coastal habitat protection and restoration activities. Aggressive land acquisition or protection initiatives (through conservation easements or other mechanisms) will be needed to ensure attainment of targets and goals for both salt marsh and upland habitats (Fig 3).

**Table 6.**
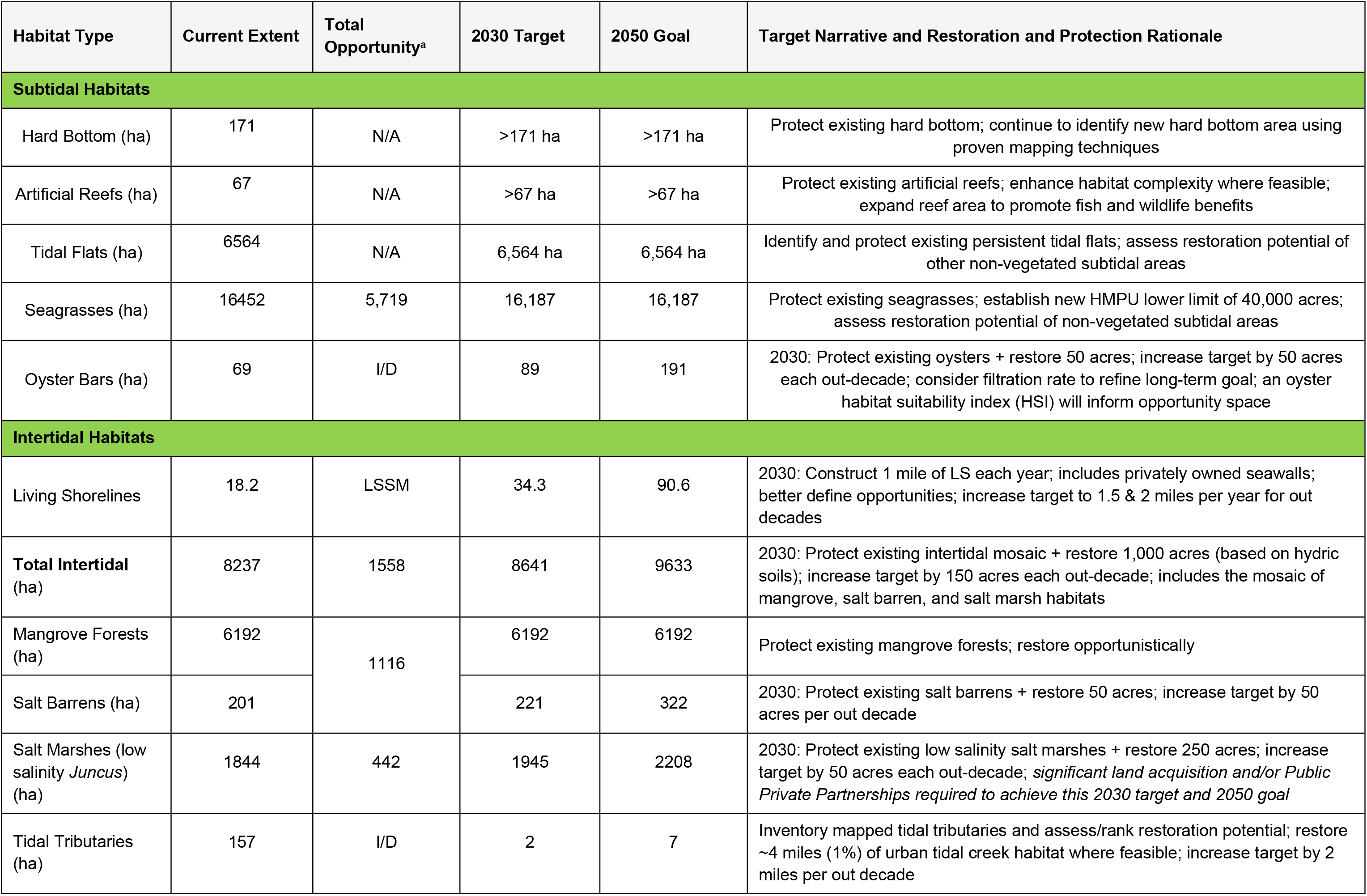
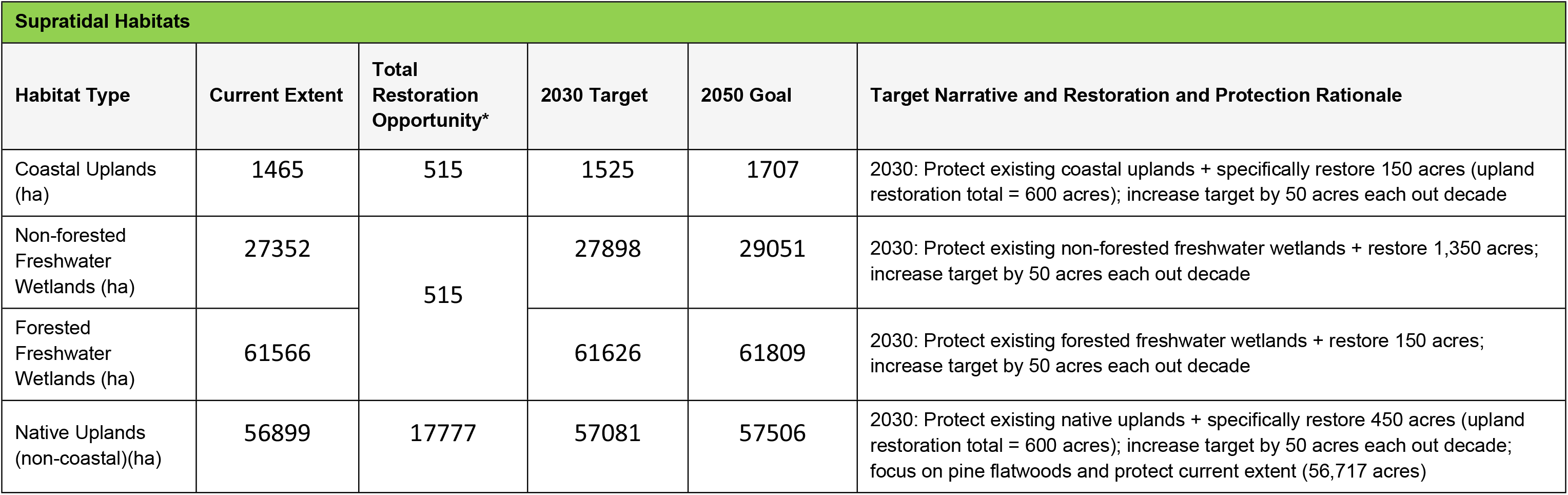
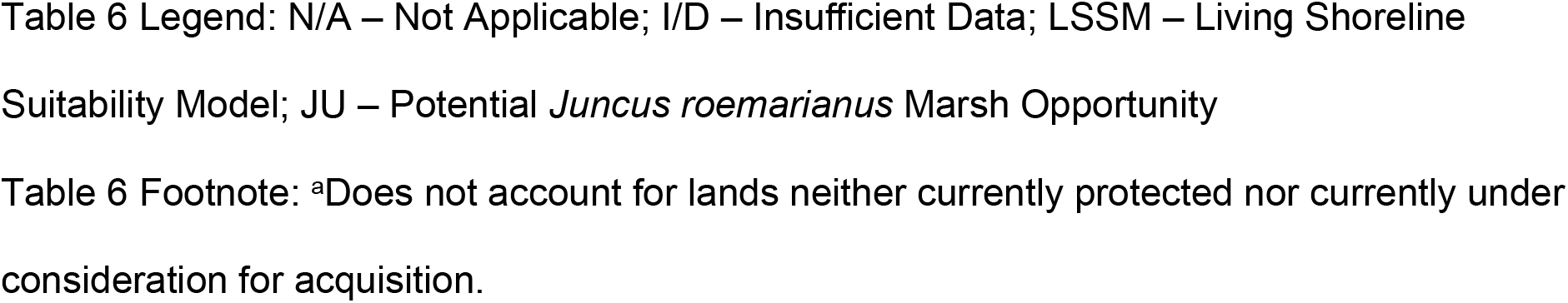
Recommended 2030 targets and 2050 goals for habitat restoration and protection in the Tampa Bay watershed.

| Habitat Type | Current Extent | Total Opportunity<sup>a</sup> | 2030 Target | 2050 Goal | Target Narrative and Restoration and Protection Rationale |
| --- | --- | --- | --- | --- | --- |
| **Subtidal Habitats** |  |  |  |  |  |
| Hard Bottom (ha) | 171 | N/A | >171 ha | >171 ha | Protect existing hard bottom; continue to identify new hard bottom area using proven mapping techniques |
| Artificial Reefs (ha) | 67 | N/A | >67 ha | >67 ha | Protect existing artificial reefs; enhance habitat complexity where feasible; expand reef area to promote fish and wildlife benefits |
| Tidal Flats (ha) | 6564 | N/A | 6,564 ha | 6,564 ha | Identify and protect existing persistent tidal flats; assess restoration potential of other non-vegetated subtidal areas |
| Seagrasses (ha) | 16452 | 5,719 | 16,187 | 16,187 | Protect existing seagrasses; establish new HMPU lower limit of 40,000 acres; assess restoration potential of non-vegetated subtidal areas |
| Oyster Bars (ha) | 69 | I/D | 89 | 191 | 2030: Protect existing oysters + restore 50 acres; increase target by 50 acres each out-decade; consider filtration rate to refine long-term goal; an oyster habitat suitability index (HSI) will inform opportunity space |
| **Intertidal Habitats** |  |  |  |  |  |
| Living Shorelines | 18.2 | LSSM | 34.3 | 90.6 | 2030: Construct 1 mile of LS each year; includes privately owned seawalls; better define opportunities; increase target to 1.5 & 2 miles per year for out decades |
| **Total Intertidal (ha)** | 8237 | 1558 | 8641 | 9633 | 2030: Protect existing intertidal mosaic + restore 1,000 acres (based on hydric soils); increase target by 150 acres each out-decade; includes the mosaic of mangrove, salt barren, and salt marsh habitats |
| Mangrove Forests (ha) | 6192 | 1116 | 6192 | 6192 | Protect existing mangrove forests; restore opportunistically |
| Salt Barrens (ha) | 201 |  | 221 | 322 | 2030: Protect existing salt barrens + restore 50 acres; increase target by 50 acres per out decade |
| Salt Marshes (low salinity *Juncus*) (ha) | 1844 | 442 | 1945 | 2208 | 2030: Protect existing low salinity salt marshes + restore 250 acres; increase target by 50 acres each out-decade; *significant land acquisition and/or Public Private Partnerships required to achieve this 2030 target and 2050 goal* |
| Tidal Tributaries (ha) | 157 | I/D | 2 | 7 | Inventory mapped tidal tributaries and assess/rank restoration potential; restore ~4 miles (1%) of urban tidal creek habitat where feasible; increase target by 2 miles per out decade |

Table 6 Legend: N/A – Not Applicable; I/D – Insufficient Data; LSSM – Living Shoreline Suitability Model; JU – Potential *Juncus roemarianus* Marsh Opportunity Table 6 Footnote: <sup>a</sup>Does not account for lands neither currently protected nor currently under consideration for acquisition.
| Supratidal Habitats |  |  |  |  |  |
| --- | --- | --- | --- | --- | --- |
| Habitat Type | Current Extent | Total Restoration Opportunity* | 2030 Target | 2050 Goal | Target Narrative and Restoration and Protection Rationale |
| Coastal Uplands (ha) | 1465 | 515 | 1525 | 1707 | 2030: Protect existing coastal uplands + specifically restore 150 acres (upland restoration total = 600 acres); increase target by 50 acres each out decade |
| Non-forested Freshwater Wetlands (ha) | 27352 | 515 | 27898 | 29051 | 2030: Protect existing non-forested freshwater wetlands + restore 1,350 acres; increase target by 50 acres each out decade |
| Forested Freshwater Wetlands (ha) | 61566 |  | 61626 | 61809 | 2030: Protect existing forested freshwater wetlands + restore 150 acres; increase target by 50 acres each out decade |
| Native Uplands (non-coastal)(ha) | 56899 | 17777 | 57081 | 57506 | 2030: Protect existing native uplands + specifically restore 450 acres (upland restoration total = 600 acres); increase target by 50 acres each out decade; focus on pine flatwoods and protect current extent (56,717 acres) |

Targets that maintain current coverage and extent (“hold-the line strategies”) were identified for habitats deemed to be at acceptable levels to support fish and wildlife resources. Evolving information such as an Oyster Habitat Suitability Index [91] and ongoing mapping exercises will be used to identify optimal locations to conduct restoration activities that help achieve targets and goals. Coordination with the establishment of state-mandated minimum flows [91] will be necessary for restoration and maintenance of low-salinity salt marsh habitats that will experience higher salinities and rapid transition to mangroves under existing sea level rise scenarios [93,94].

## Discussion

Because of multiple stressors such as encroaching development and climate change, habitat protection and restoration priorities should be tempered and “reality tested” by what is feasible now and what may be possible in the future. Many native and potentially restorable habitats are limited, and there will always be restrictions on the financial resources that can be dedicated to promote public conservation land acquisition and habitat restoration activities. However, there continues to be recognition of the importance of these efforts to protect biodiversity and ecosystem services [95,96]

This replicable, regional and watershed-based methodology for setting restoration targets and goals provides a systematic attempt to identify habitat protection and restoration targets that are based on what is actually achievable within the limitations of a continuously urbanizing coastal setting. It focuses on existing opportunities for all habitat types, and what is realistically possible in the future, rather than attempting to mimic previous ecological conditions, that are more or less impractical to replicate within the current setting [34].

### Habitat trends

When viewed as a whole, the most significant and meaningful trends in the habitats of interest over the periods of record examined include: 1) the 75 percent gain in seagrasses since 1988; 2) the slight gains in emergent tidal wetlands (10%) and freshwater wetlands (2%) since 1990; and 3) the 39% loss in native upland habitats since 1990.

The intertidal zone in Tampa Bay is currently experiencing dynamic change, driven by sea level rise and climate change, whereby mangrove forests are outcompeting salt marshes and salt barrens for the available niche space. Without increasing the total area of the intertidal zone, restoring a greater coverage of mangroves would reduce the niche space available for salt marshes and salt barrens. This phenomenon has been observed throughout the Gulf of Mexico, and has been attributed to both climate change (e.g., fewer freeze events) and sea level rise [58].

The observed gains in wetlands (Table 3b) are likely a reflection of: 1) the effectiveness of state and federal wetland regulatory programs; and 2) the cumulative gains resulting from, primarily, publicly-funded habitat restoration projects (Table 4). Minor gains in some emergent tidal wetlands (e.g., salt barrens) may also be a reflection of the landward expansion of the complex suite of these habitats associated with climate change and sea level rise. Gains in vegetated non-forested freshwater wetlands are related to the clearing of forested wetlands followed by the creation of herbaceous mitigation areas and stormwater systems.

The decrease in native uplands (Table 3b) is the result of continued development in the Tampa Bay watershed, combined with the lack of regulatory protection of native uplands. Attaining upland habitat targets and goals will require coordinated and concerted restoration of these habitats on existing conservation lands, as well as new conservation lands to offset the continued loss of these habitats to development. Further, amendments to existing planning, zoning and land development policies or regulations will need to be examined, if the region is to fully embrace the aspirational objectives for these habitats. While federal and state regulations related to listed species management impart some protection to certain rare upland habitats -- such as scrub jay (*Aphelocoma coerulescens*) habitat -- common and historically abundant native habitats, such as pine flatwoods, have been left largely unprotected. Substantial opportunities also exist for upland restoration on reclaimed mined lands within the watershed. However, unless local governments in the Tampa Bay watershed improve local protections for native uplands, such as strengthening language within comprehensive plans and development ordinances, this trend will likely continue into the future [95,98].

### Habitat restoration

Increases observed in tidal and freshwater wetlands are primarily due to publicly-funded habitat restoration projects, state and federal wetland regulatory programs, and to a lesser extent, regulatory mitigation. While restoration activities date to 1971 in the Tampa Bay region, few projects were completed prior to 1990, and from 1990-2010, an annual mean of 68 ha of habitat was restored. Over the past decade (2010-2019), the rate of restoration project completion and investment has increased to over 81 ha/yr. Assuming that funding levels remain in the same range as the past decade, this annual mean can be used to set reasonable limits on restoration potential and targets.

While existing development areas are not considered feasible for major habitat restoration activities at this time, there are many opportunities to enhance and restore habitat functions and improve coastal resilience in urbanized locations. Examples include the construction of living shorelines, placement of submerged habitat modules along developed urban shorelines and seawalls, and creation of backyard habitats. Tidal tributary restoration could also entail improvements in hydrology, such as removal of salinity barriers and filling of dredged channel sections prone to low dissolved oxygen levels [61,64].

Four major types of disturbed sites around the Tampa Bay coastline have been identified as priority estuarine habitat restoration sites by stakeholders over the past two decades, including dredged holes, filled and spoil disposal areas, abandoned aquaculture ponds; and coastal borrow pits and stormwater ponds.

There is a general consensus among restoration practitioners and natural resource managers that habitat restoration and management is most cost-effective on publicly-owned conservation lands and provides long-term benefits. Furthermore, given current development trends in the Tampa Bay watershed, public acquisition of remaining critical lands (e.g., coastal uplands, river floodplain wetlands) is a high priority, and some restoration targets (e.g., salt marshes) will not be feasible without additional public acquisition or public-private partnerships [95,99]. Therefore, varied approaches to leverage resources, including traditional grants, partner funding, and use of volunteers for habitat restoration are recommended to maximize the potential for successful target and goal achievement.

### Rolling easements, mitigation and restoration consortium

As discussed, land acquisition for coastal habitat restoration must prioritize adjacent low-lying coastal uplands to serve as buffers to accommodate future landward migration of tidal wetlands in response to sea level rise [93]. Where public acquisition is not possible, other conservation mechanisms need to be explored. Coastal setbacks, buffers, or public easements are traditionally used to restrict development within a given distance from the shoreline. A rolling easement is a dynamic mechanism that “rolls” landward as sea levels rise and cause tidal encroachments onto low-lying coastal uplands [16]. The application of rolling easements in Tampa Bay could disincentivize more intense urban development (e.g., discourage up-zoning) of low-lying coastal uplands that may be currently in less intense agricultural or recreational (e.g., golf courses) land uses. Under a rolling easement, landowners would be able to maintain current economic uses, while “reserving” such lands to accommodate tidal wetland migration with future advancing sea level rise.

Wetland impacts and associated compensatory mitigation projects authorized under wetland regulatory programs have historically been conducted independent of watershed-level planning and monitoring processes. This disconnect has contributed to fragmented implementation and inconsistent compliance monitoring of mitigation projects, as well as historically poor documentation of wetland losses and gains in the Tampa Bay watershed. However, if properly focused and comprehensively coordinated, compensatory mitigation activities could significantly contribute to the attainment of wetland habitat restoration goals and targets for the Tampa Bay estuarine system and its contributing watershed.

To better coordinate future activities, stakeholders are beginning to pledge the formation of a public-private partnership that would link regulatory (compensatory mitigation) and resource management objectives forpublicly funded habitat enhancement, restoration, and establishment programs in the watershed. Creation of a “Habitat Management Consortium” would be expected to optimize and improve the cost-effectiveness of habitat protection, restoration and mitigation activities in the watershed, and bind commitments to attaining the aspirational targets and goals for all critical coastal habitats of the Tampa Bay estuary.

## Conclusion

The establishment of habitat restoration targets and goals considering climate change, development trends, land availability, and past restoration activities, expands the restoration palette to a more comprehensive list of habitats within the system. If successfully implemented, the 2050 goals would total over 4,000 ha of habitat restoration throughout the Tampa Bay watershed. Land acquisition will be an important component of attainment of some targets and goals, including the salt marsh restoration target for the first ten years. Land acquisition will also provide new opportunities for socioeconomic benefits, given that these projects often have a public recreation component.

Our new approach will continue to engage multiple partner agencies, non-governmental organizations, and private citizens in the successful implementation of the restoration plan. Emphases will include recognition of land types particularly vulnerable to climate change or development stressors and these needs will be communicated to restoration partners. Consistent education, targeted funding opportunities, and reporting will also ensure that these newly established targets and goals lead to successful restoration projects, land acquisition, and enhanced ecosystem services.

## Acknowledgements

The TBEP Technical Advisory Committee reviewed drafts of the original reports that are the basis of this paper, and their comments and insight are gratefully recognized.

